# Defense systems and horizontal gene transfer in bacteria

**DOI:** 10.1101/2024.02.09.579689

**Authors:** Roman Kogay, Yuri I. Wolf, Eugene V. Koonin

## Abstract

Horizontal gene transfer (HGT) is a fundamental process in the evolution of prokaryotes, making major contributions to diversification and adaptation. Typically, HGT is facilitated by mobile genetic elements (MGEs), such as conjugative plasmids and phages that generally impose fitness costs on their hosts. However, a substantial fraction of bacterial genes is involved in defense mechanisms that limit the propagation of MGEs, raising the possibility that they can actively restrict HGT. Here we examine whether defense systems curb HGT by exploring the connections between HGT rate and the presence of 73 defense systems in 12 bacterial species. We found that only 6 defense systems, 3 of which are different CRISPR-Cas subtypes, are associated with the reduced gene gain rate on the scale of species evolution. The hosts of such defense systems tend to have a smaller pangenome size and harbor fewer phage-related genes compared to genomes lacking these systems, suggesting that these defense mechanisms inhibit HGT by limiting the integration of prophages. We hypothesize that restriction of HGT by defense systems is species-specific and depends on various ecological and genetic factors, including the burden of MGEs and fitness effect of HGT in bacterial populations.

## Introduction

Bacterial viruses, known as bacteriophages (phages, for short), are the most abundant entities in the biosphere (<u>Keen, 2015</u>). They regularly attack and predate on bacterial populations across different ecological settings with the estimated rate of infection per second in oceans alone is on the order of 10^23^ (<u>Suttle, 2007</u>; <u>Mushegian, 2020</u>). To counteract phages and other parasitic mobile elements, bacteria evolved a wide range of defense systems with various molecular mechanisms of action (<u>Doron et al., 2018</u>; <u>Gao et al., 2020</u>; <u>Bernheim et al., 2021</u>; <u>Millman et al., 2022</u>; <u>Georjon and Bernheim, 2023</u>). These include CRISPR-Cas systems, which provide adaptive immunity by storing information about past encounters with MGE (<u>Makarova et al., 2020</u>), restriction-modification (RM) systems that degrade foreign genetic material based on specific molecular patterns (<u>Wilson, 1991</u>), abortive infection mechanisms that limit the spread of phages in the bacterial population by inducing suicide of infected cells (<u>Lopatina et al., 2020</u>), and multiple others. Individual bacterial genomes typically encode several diverse defense systems, and the repertoire of defense mechanisms can differ even among closely related strains (<u>Bernheim and Sorek, 2020</u>; <u>Tesson et al., 2022</u>). Consequently, defense systems demonstrate high mobility, with high rates of gene gain and loss on a short evolutionary scale (<u>Makarova et al., 2013</u>; <u>Puigbo et al., 2017</u>).

Although defense systems are essential for protection against phages, and to a lesser extent, against other invasive MGEs, such as integrative conjugative elements (ICE) and plasmids, they also come with associated fitness costs to the hosts. One form of such costs is impediment to lysogenic conversion and gain of beneficial genes that reside in MGEs. These include genes that equip bacteria with the capability to adapt to different ecological niches (<u>Kelleher et al., 2017</u>; <u>Davray et al., 2021</u>; <u>Kieft et al., 2021</u>), and resist environmental stress (<u>Lopatkin et al., 2017</u>; <u>Jahn et al., 2019</u>). For example, the presence of CRISPR-Cas system in *Enterococcus faecalis* shows a significant inverse correlation with the resistance to different antibiotics (<u>Palmer and Gilmore, 2010</u>). Moreover, multiple experimental studies have demonstrated the capability of CRISPR-Cas systems to limit horizontal gene transfer (HGT) (<u>Marraffini and Sontheimer, 2008</u>; <u>Bikard et al., 2012</u>). However, broader comparative genomic analyses yielded conflicting conclusions on the inhibition of HGT by CRISPR-Cas on a larger evolutionary scale (<u>Gophna et al., 2015</u>; <u>Shehreen et al., 2019</u>; <u>Wheatley and MacLean, 2021</u>). Furthermore, potential interference of other defense systems with HGT has not be comprehensively analyzed.

In this work, we examined the association between the presence of various defense systems and the rates of gene gain in a set of 12 bacterial species. Our results reveal significant association with increased gene gain rate for 15 defense systems, whereas 6 systems were found to be significantly associated with reduced gene gain rates. However, we show that for the 15 defense systems associated with increased gene gain rates, this signal is likely a byproduct of their location within large MGEs. Conversely, 3 of the 6 defense systems that are negatively correlated with the gene gain are CRISPR-Cas variants that tend to inhibit gene gain by reducing prophage integration.

## Results

### Only a few defense systems are associated with reduced gene gain rate

We analyzed a dereplicated set of 2,546 complete genomes from 12 bacterial species. Each species was represented by a minimum of 118 genomes, with the count ranging from 118 for *Streptococcus pyogenes* to 430 for *Escherichia coli* (**Supplementary Table S1**). Additionally, each species was predicted to possess at least two defense systems encoded in at least 20% of the genomes, but not more than in 80%.

To estimate the gene gain rate for each species, we mapped the ancestral state of every orthologous group using GLOOME (<u>Cohen et al., 2010</u>) onto the individual species trees, which were computed from the concatenated alignments of the core genes (see **Experimental Procedures** for the details). GLOOME estimates the probability of the presence of each orthologous groups in the ancestral nodes, taking a value from 0 to 1. We then calculated changes in the probabilities across all individual ancestral states from the root to the tips of phylogenetic trees. The positive differences were retained and summed up across all orthologous groups to obtain the overall gene gain estimation for every branch in the phylogenies.

Additionally, the combined gene gain estimates were normalized by their respective branch lengths. For the defense systems, we also conducted the ancestral state reconstruction using GLOOME (<u>Cohen et al., 2010</u>), and denoted the state of every branch in phylogenetic trees as the average of the states between its ancestral and descendant nodes. To determine whether there was a connection between individual defense systems and gene gain rate, we calculated the Spearman’s correlation coefficient between them (**Supplementary Figure S1**). The computational pipeline developed for this analysis is shown in **Figure 1**.

**Figure 1.**
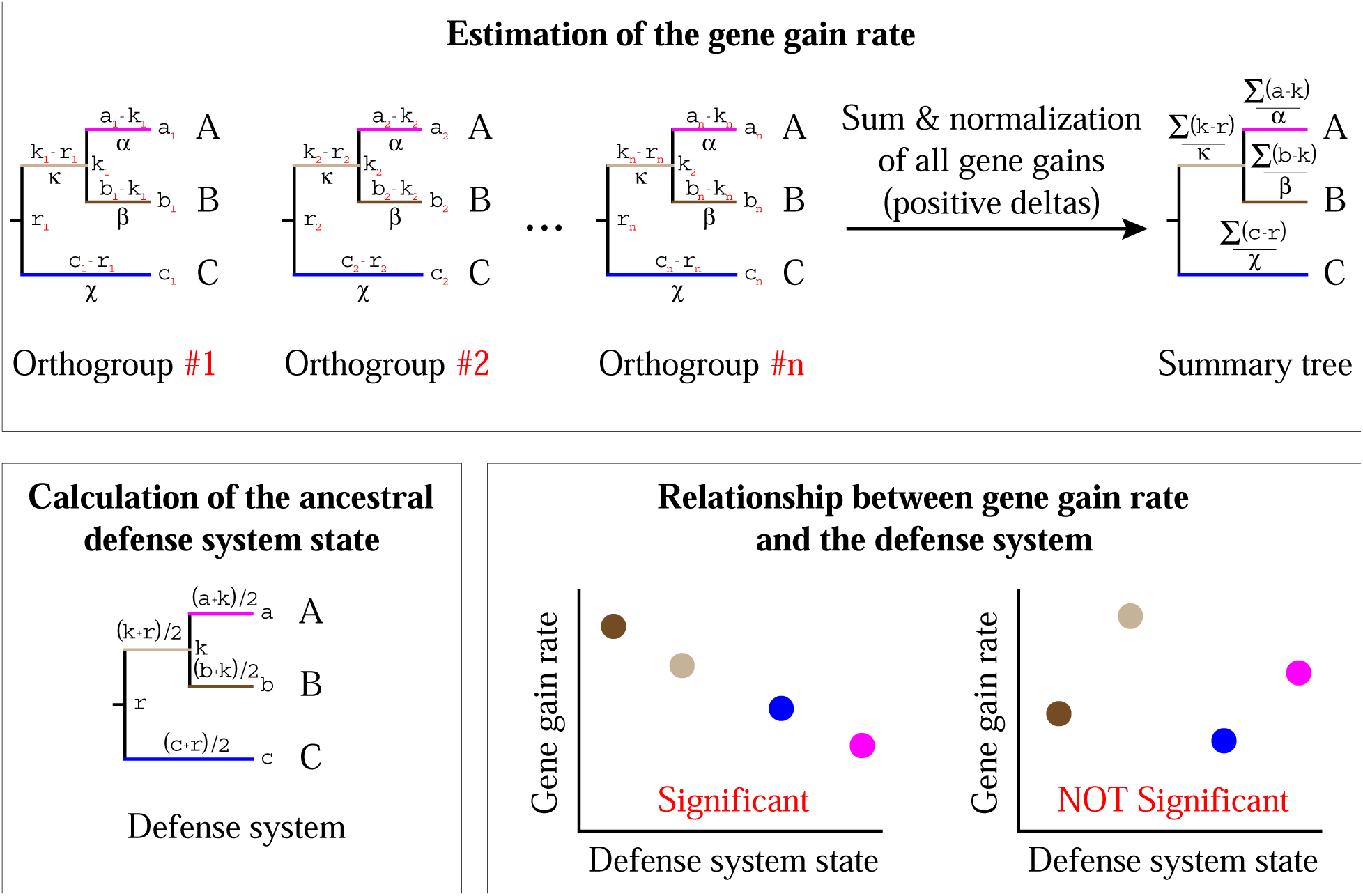
The pipeline used to calculate the association between the presence of defense systems and gene gain rate. In the initial step, the gene gain rate is calculated for each branch of the phylogenetic tree. Subsequently, the probabilistic state of a defense system is mapped onto the phylogenetic tree. In the final step, the Spearman’s correlation coefficient between the obtained estimates is computed.

Of the 73 analyzed defense systems, 6 were found to be significantly associated with a reduced gene gain rate (Spearman’s rho < –0.3; permutation test p-value < 0.0001), whereas 15 were significantly associated with an increased gene gain rate (Spearman’s rho > 0.3; permutation test p-value < 0.0001) (**Figure 2**, **Supplementary Table S2**). Notably, among the 6 defense system associated with a reduced gene gain rate, 3 were CRISPR-Cas variants.

**Figure 2.**
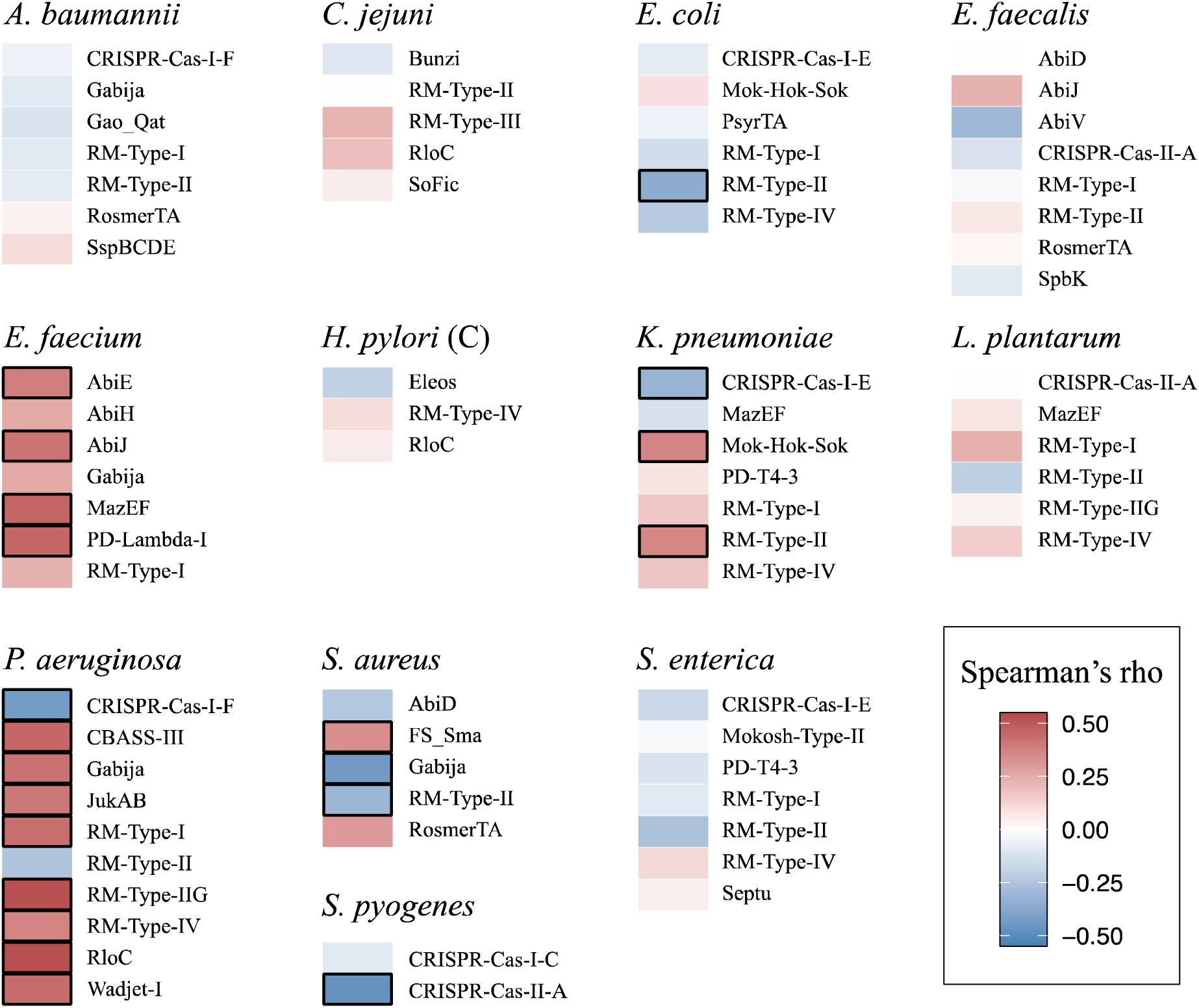
Heatmap of the correlation coefficients between defense systems and gene gain rate. Positive correlation coefficients are shown with red hues, and negative correlation coefficients are shown with blue hues. Significant Associations (|r_S_ | > 0.3) are highlighted with black rectangles.

Given that HGT plays the key role in the expansion of the accessory genome in bacteria (<u>Ochman et al., 2000</u>; <u>Nowell et al., 2014</u>; <u>Props et al., 2019</u>), we hypothesized that defense systems associated with a reduced gene gain rate would be present in pangenomes of a relatively smaller size, and vice versa. To test this hypothesis, we counted the number of protein coding genes and conducted pairwise comparisons of the genomes containing and lacking the respective defense systems. Indeed, we found that defense systems that are associated with a reduced gene gain rate tend to be present in genomes that encode fewer proteins, and in 3 of the cases, this difference was statistically significant (Mann-Whitney U test, p-value < 0.05) (**Figure 3**). To account for the phylogenetic signal, we also performed phylogenetic logistic regression (<u>Ives and Garland, 2010</u>), and in all 3 cases, the inverse relationship remained significant (p-value < 0.05). In contrast, we expected that genomes carrying the defense systems associated with an elevated gene gain rate would contain more protein-coding genes. Indeed, in each of the 15 such cases, genomes that harbored these defense systems had significantly larger pangenome size compared to genomes lacking them (Mann-Whitney U test, p-value < 0.05) (**Supplementary Figure S2**), and the statistical significance persisted after applying the phylogenetic correction (p-value < 0.05).

**Figure 3.**
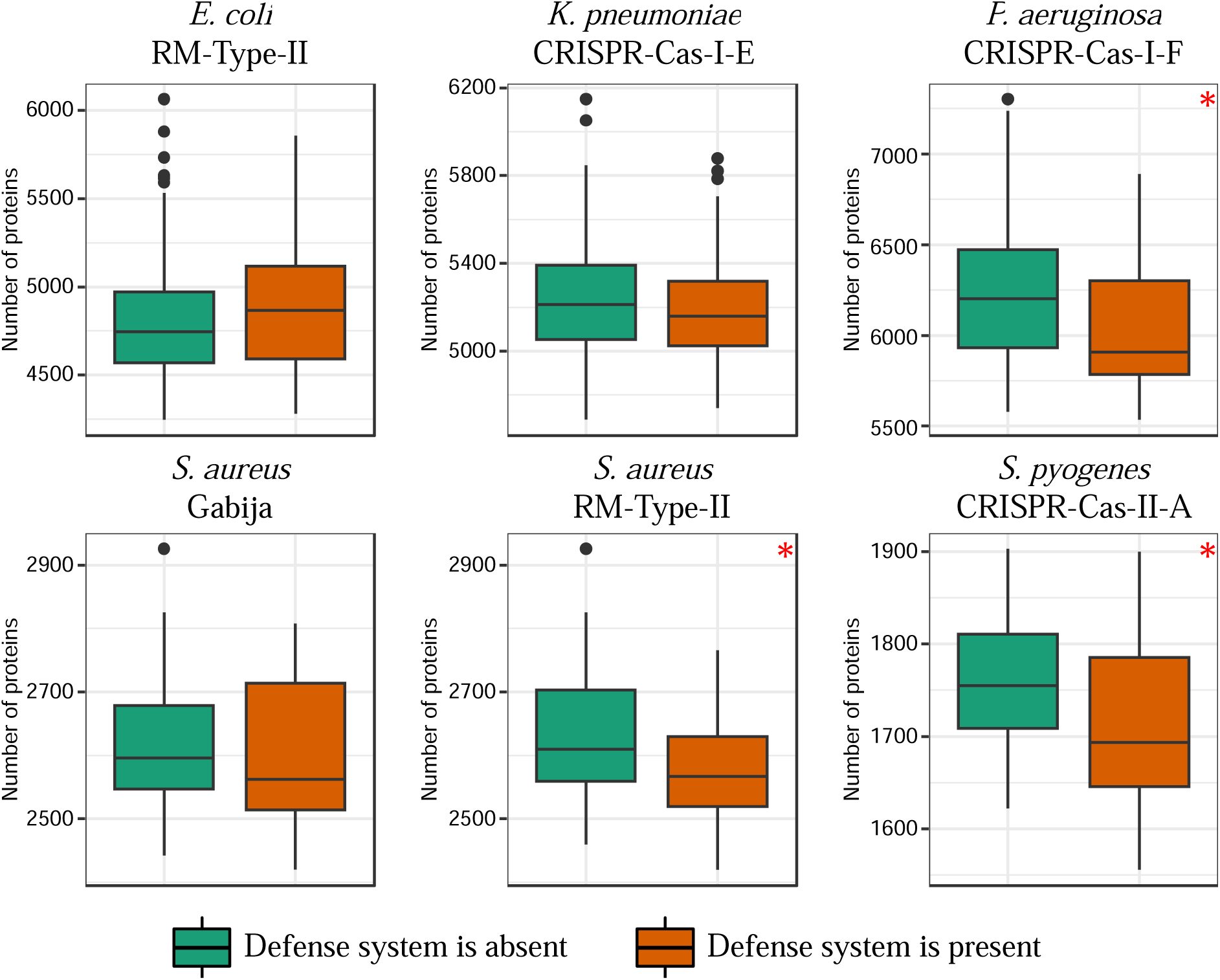
Relationship between the 6 defense systems associated with a reduced gene gain rate and the number of protein-coding genes in genomes. Boxplots show median values that are bound by the first and third quartiles. Whiskers extended outside boxplots are within the 1.5 * interquartile range. Black dots outside whiskers denote outliers. Red asterisks show significant difference between boxplots (Mann-Whitney U test, p-value < 0.05).

We observed that 3 CRISPR-Cas systems, including CRISPR-Cas-I-F in *Pseudomonas aeruginosa*, were significantly associated with a reduced gene gain rate, corroborating previous work (<u>Wheatley and MacLean, 2021</u>). By contrast, 6 CRISPR-Cas systems, including CRISPR-Cas-I-E in *Escherichia coli* and *Salmonella enterica*, exhibited no significant link with the gene gain rate, also in agreement with previous findings (<u>Gophna et al., 2015</u>; <u>Shariat et al., 2015</u>; <u>Xue and Sashital, 2019</u>) (**Figure 2)**. As a proxy for the CRISPR-Cas activity, we calculated spacers turnover rate in the respective species (see **Experimental Procedures** for the details). Active CRISPR-Cas systems tend to continuously acquire new spacers (<u>Paez-Espino et al., 2013</u>), whereas old spacers are prone to be lost due to the inactivity and pronounced deletion bias in bacteria (<u>Mira et al., 2001</u>). Thus, more active CRISPR-Cas systems are likely to have a higher spacer turnover rate. Indeed, although the signal was not particularly robust, we found that species where CRISPR-Cas was associated with reduced gene gain rate tended to have a higher spacer turnover rate (**Supplementary Figure S3**) (**Supplementary Table S3**).

### Defense systems reduce the gene gain rate by limiting acquisition of phage-like genes

Given that the primary role of defense systems in bacteria is to limit the propagation of a phage infection and, to a lesser extent, the spread of ICEs and plasmids, we hypothesized that defense systems that are associated with a reduced gene gain rate actively hinder the spread of MGEs. Consequently, genomes that encompass such defense systems can be expected to harbor fewer phage-related genes than other genomes. Indeed, our findings show that the genomes carrying 4 of the 6 defense systems that are associated with a reduced gene gain rate, including all 3 CRISPR-Cas systems, contained a significantly lower number of phage-related genes compared to genomes that lack such defense systems (Mann-Whitney U test, p-value <0.05) (**Figure 4**). These results are also supported by the phylogenetic logistic regression (p-value < 0.05). We conjectured that these CRISPR-Cas systems play a direct role in restricting the acquisition of phage-related genes, that is, curtail prophage integration. To test this hypothesis, we investigated whether the spacers in these CRISPR-Cas systems targeted phage genes present in the respective species. For that purpose, we pooled all detected phage-related genes encoded in genomes of the respective species and found that spacers target a range of 1.48% to 6.71% of all phage-related genes (**Supplementary Figure S4A**). Notably, phage-related genes targeted by spacers are predominantly found in genomes lacking the 3 CRISPR-Cas systems associated with a reduced gene gain rate (**Supplementary Figure S4B**). These observations are compatible with the possibility that these CRISPR-Cas systems actively inhibit the acquisition of prophages.

**Figure 4.**
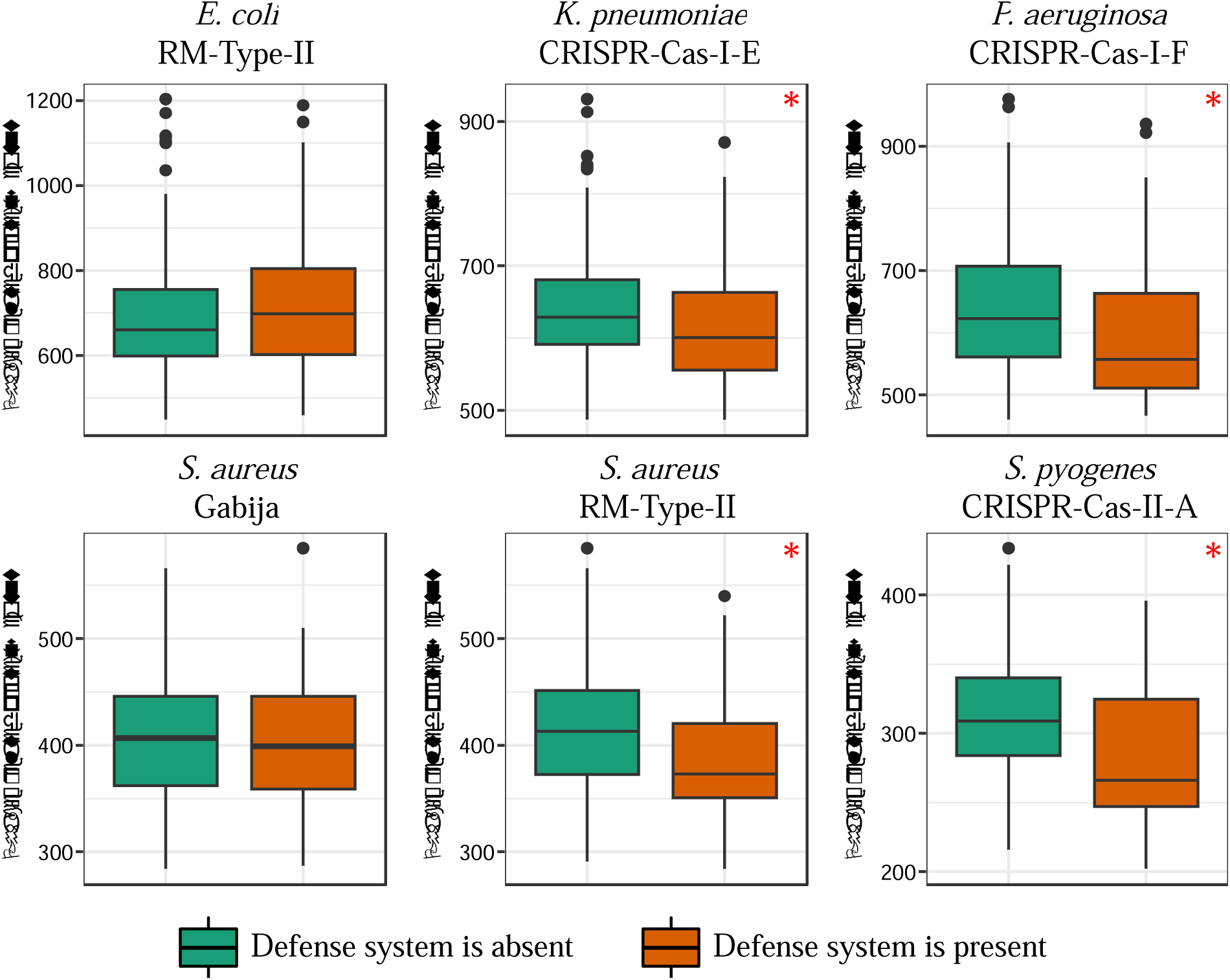
Relationship between the 6 defense systems associated with a reduced gene gain rate and the number of phage-related genes in genomes. Boxplots show median values that are bound by the first and third quartiles. Whiskers extended outside boxplots are within the 1.5 * interquartile range. Black dots outside whiskers denote outliers. Red asterisks show significant differences between boxplots (Mann-Whitney U test, p-value < 0.05).

We also analyzed the number of phage-related genes in genomes that harbored defense systems associated with increased gene gain rate. The results indicated that genomes carrying such defense systems contained more phage-related genes than genomes without these systems, and in all 15 cases, that difference was statistically significant (**Supplementary Figure S5**), as supported by phylogenetic logistic regression (p-value < 0.05).

### Defense systems are linked to mobile genetic elements

Because defense systems tend to be colocalized with MGEs in prokaryotic genomes (<u>Makarova et al., 2011</u>), we hypothesized that the connection between some defense systems and an increased gene gain rate is a byproduct of horizontal transfer of large ICEs and plasmids carrying these defense systems. To address this possibility, we conducted phylogenetic profiling of 47 defense systems whose constituent genes form orthologous groups across the genomes in examined species. To minimize false positive associations due to the phylogenetic signal, we discarded clade-specific defense systems and the corresponding groups of orthologous genes (see **Experimental Procedures** for the details). For every pair of orthologous groups, we calculated the observed and expected counts of occurrence in a genome, and evaluated the significance using the Bonferroni-corrected binomial test (<u>Whelan et al., 2020</u>). If defense systems and co-occurring orthologous groups shared evolutionary and functional connections, they would be expected to co-localize in bacterial genomes. We calculated median genomic distance between defense systems and co-occurring orthologous gene groups and compared them with the distribution expected by chance that was generated by random sampling of the orthologous gene groups encoded in the main chromosome. Indeed, we found that of the 782 identified associations, 725 orthologous gene groups substantially colocalized with the defense systems (p-value < 0.1).

Furthermore, our findings indicate that defense systems associated with the increased gene gain rate co-occurred with significantly more orthologous gene groups compared to other defense systems (Mann-Whitney U test, p-value < 0.05). A closer examination of the orthologous gene groups over-represented in the genomic neighborhoods of defense systems suggested that these neighborhoods predominantly represented ICEs and large extrachromosomal plasmids, as exemplified by JukAB, RM-Type-IV and Wadjet systems in *Pseudomonas aeruginosa* (**Figure 5A**). Consequently, the association between these defense systems and an increased gene gain rate could be attributed to horizontal movement of MGEs that harbor such defense systems along with the co-occurring genes. Moreover, in agreement with prior findings (<u>Makarova et al., 2011</u>; <u>Botelho, 2023</u>), 19 of the 26 defense systems with at least one associated orthologous gene group were linked to genes typical of MGEs including phages, ICEs, transposons and integrons. Except for CRISPR-Cas-I-E in *Klebsiella pneumoniae*, CRISPR-Cas systems shared no phylogenetic co-occurrence with other orthologous gene groups (**Figure 5B**).

**Figure 5.**
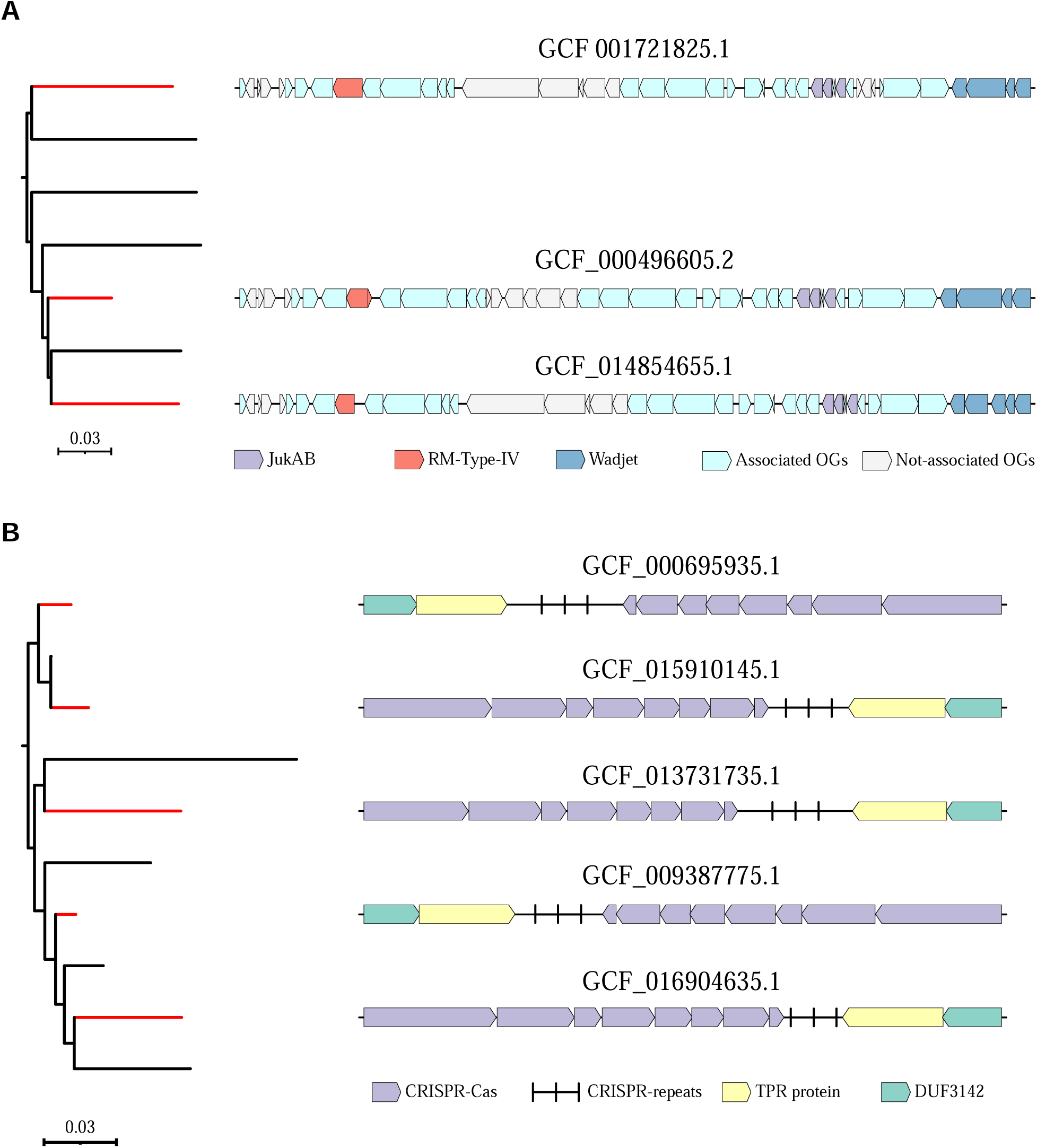
Gene neighborhoods of (A) JukAB, RM-Type IV and Wadjet in *Pseudomonas aeruginosa*, and (B) CRISPR-Cas-I-E in *Klebsiella pneumoniae* in representative genomes. Phylogenetic trees are scaled with the number of substitutions per site, and red branches denote clades that contain discussed defense systems.

## Discussion

Experiments have clearly demonstrated that bacterial defense systems can actively limit propagation of MGE such as phages and plasmids under laboratory conditions (<u>Marraffini and Sontheimer, 2008</u>; <u>Dupuis et al., 2013</u>; <u>Deep et al., 2022</u>). However, whether on the larger evolutionary scale these systems provide any substantial barrier to the horizontal gene acquisition, remains a conflicting topic (<u>Gophna et al., 2015</u>; <u>Wheatley and MacLean, 2021</u>). Our results indicate that the majority of the defense systems are not associated with either an increased or a decreased rate of gene acquisition. However, a minority of the defense systems are associated with an elevated gene gain rate, and an even a smaller subset is associated with a reduced gene gain rate.

While certain CRISPR-Cas systems, Gabija and RM-Type II are associated with a reduced gene gain rate in some species, in other species, they do not exhibit such association. Thus, the effects of defense systems on HGT appear to be strongly lineage-specific and depend on additional factors. Integrated phages and plasmids can impose various metabolic and other fitness costs on their hosts contingent upon ecological and genetic contexts (<u>Alonso-Del Valle et al., 2021</u>; <u>Rendueles et al., 2023</u>). Hence, there is likely a differential selection pressure on same types of defense systems in different species to hinder the spread of MGEs, depending on the associated costs of the latter. Perhaps, more generally and more importantly, given that HGT is the major route of acquisition of novel traits by bacteria, the active restriction of the gene flow can compromise bacterial adaptation to diverse, fluctuating environmental conditions (<u>Woods et al., 2020</u>; <u>Arnold et al., 2022</u>). In this scenario, the costs associated with defense systems can outweigh their benefits, leading to a reduction in their activity or even complete inactivation and subsequent loss. Furthermore, the extent of HGT varies among different bacterial species and strains depending on the ecological conditions (<u>Smillie et al., 2011</u>; <u>Groussin et al., 2021</u>). For example, niche specialists that occupy stable environmental habitats tend to have closed pangenomes and lower genetic diversity relative to generalists with open pangenomes (<u>Brockhurst et al., 2019</u>). As a result, for some bacteria, ecological barriers to HGT could be so pronounced that the restriction of HGT by defense systems becomes limited and could be neither statistically significant nor biologically relevant.

Bacteria rely on different molecular strategies, including innate immunity, adaptive immunity and abortive infection to combat infection by phages and restrict the invasion of other costly MGEs, such as conjugative plasmids (<u>Makarova et al., 2021</u>). Such multilayered defense organization provide cells with an enhanced capability to withstand assaults by diverse MGEs. Abortive infection is typically used by bacteria as a last-resort defense strategy during the final stages of phage reproduction when the cell lysis becomes imminent (<u>Lopatina et al., 2020</u>; <u>Rousset and Sorek, 2023</u>). Therefore, integration of prophages or uptake of conjugative plasmids are not likely to trigger this type of immune response, and consequently, abortive infection appears unlikely to substantially interfere with HGT facilitated by phages and other MGEs.

Indeed, among the 6 defense systems we found to be associated with a reduced gene gain rate, none is known to be involved in abortive infection response. By contrast, 3 of these defense systems are CRISPR-Cas variants from *Pseudomonas aeruginosa*, *Klebsiella pneumoniae* and *Streptococcus pyogenes* that appear to restrict gene gains, primarily, by interfering with prophage integration. On the other hand, we found no evidence that CRISPR-Cas-I-E in *Escherichia coli* and *Salmonella enterica* impacts gene acquisition on the species-level, which is consistent with prior work demonstrating the low activity of these systems (<u>Westra et al., 2010</u>; <u>Shariat et al., 2015</u>).

Our observations of a positive association with some defense systems with HGT rate seem to be explained away by their hijacking of MGE for. Nevertheless, bona fide stimulation of HGT by defense systems cannot be ruled out. For example, in *Petrobacterium atrosepticum*, CRISPR-Cas-I-F can promote HGT through generalized transduction by boosting the survival rate of cells that receive transduced genetic material during infection, while the defense system inhibits lytic phages (<u>Watson et al., 2018</u>). Such mechanism could represent an adaptation to facilitate the gene flow while maintaining an active defense system that deters deleterious MGEs. Consequently, the interplay of population level dynamics among various MGEs and bacteria that harbor defense systems can determine the varying effects of defense systems on HGT.

## Experimental Procedures

### Generation of the dataset

All bacterial assemblies with the ‘complete genome’ assembly level were downloaded from the RefSeq database (assessed October 20, 2023) (<u>O’Leary et al., 2016</u>). 30,177 assemblies with the completeness greater than 90% and contamination less than 5%, as determined by CheckM, were retained for subsequent analyses (<u>Parks et al., 2015</u>). The taxonomy of each retained assembly was determined either by assessing the GTDB taxonomy database or via *de novo* prediction using the gtdb-tk v2.3.2 (<u>Parks et al., 2018</u>; <u>Chaumeil et al., 2019</u>). Average Nucleotide Identity (ANI) values were calculated using the fastANI v1.33 (<u>Jain et al., 2018</u>) and genomes were dereplicated using the ANI cutoff of 99.9% via a single-linkage clustering algorithm (due to the substantially larger number of available genomes, for *E. coli* and *K. pneumoniae,* the ANI threshold was reduced to 99.5% and 99.8%, respectively). Fifteen species represented by at least 100 dereplicated genomes were retained.

Genomes from the retained species were annotated using Prokka v1.14.6 with default parameters (<u>Seemann, 2014</u>). In each retained genome, defense systems were predicted using DefenseFinder (last updated November, 2023) (<u>Abby et al., 2014</u>; <u>Tesson et al., 2022</u>). Twelve species that encoded a minimum of two defense systems found in at least 20% of genomes, but not more than in 80%, were retained for subsequent analyses (**Supplementary Table S1**).

### Identification of orthologous gene groups and reconstruction of reference phylogenetic trees

For each retained species, orthologous groups were predicted using the Panaroo v1.3.4 with the sensitive mode (<u>Tonkin-Hill et al., 2020</u>). Protein sequences encoded by genes from orthologous groups that are found in at least 95% of the genomes were aligned using MAFFT v7.520 (<u>Katoh and Standley, 2013</u>). Phylogenetically informative alignments were filtered using BMGE (Block Mapping and Gathering with Entropy) as implemented in Panaroo (<u>Criscuolo and Gribaldo, 2010</u>). The retained alignments were concatenated into a single superalignment. The superalignments were trimmed using ClipKIT v2.1.1 to keep only parsimony-informative sites (<u>Steenwyk et al., 2020</u>), and phylogenetic trees were reconstructed using IQ-Tree v2.2.5 with GTR+I+G model (<u>Minh et al., 2020</u>). The reconstructed phylogenetic trees were rooted using the minimum ancestral deviation method (<u>Tria et al., 2017</u>). Phylogenetic trees were visualized in iToL v6 (<u>Letunic and Bork, 2021</u>).

### Correlation between the gene gain rate and defense system presence

The presence-absence matrix of orthologous gene groups was transformed into binary sequences in FASTA format that was used as input for GLOOME (<u>Cohen et al., 2010</u>). The reconstructed ancestral states and data for the original taxa were mapped onto the reference phylogenetic trees. For each orthologous group, the changes for every branch were calculated by subtracting each ancestral state from the descendant state and keeping the results only if the difference was positive. All retained changes across all orthologous groups were combined into a single number and normalized by their respective branch lengths. In a similar manner, the ancestral states were reconstructed for each defense system, and for every branch in the phylogenetic tree, the state of each defense system was represented as an average between its ancestral and descendant states. Then, the Spearman’s correlation coefficient was calculated for the gene gain rate and the presence of each defense system. To calculate the p-value, values of the gene gain rate and the state of defense systems were randomly shuffled, and the Spearman’s correlation coefficient was calculated for the shuffled dataset. This procedure was repeated 10,000 times to generate a null distribution. To minimize the false positives, the association between gene gain rate and defense system was deemed significant if the absolute value of Spearman’s correlation coefficient was greater than 0.3, and the p-value was less than 0.0001.

### Computation of the spacers turnover rate

Genomes encompassing CRISPR-Cas systems were searched for spacers using CRISPRIdentify (<u>Mitrofanov et al., 2021</u>). Only “bona-fide” and “possible” candidates were retained for further analyses. Additionally, only spacers located in the same genetic neighborhood (within a 10,000 bp range) as the analyzed CRISPR-Cas systems were kept. For each species, an all-vs-all search of spacers was performed using MMseqs2 under the high sensitivity mode and the query and subject overlap of at least 80% (e-value < 0.01) (<u>Steinegger and Soding, 2017</u>). For every pair of taxa with the pairwise phylogenetic distance of less than 1, the spacer turnover rate was calculated as:

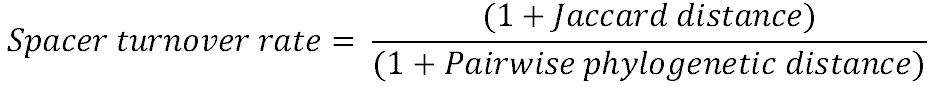

### Prediction of phage-related genes and search for corresponding spacers

HMM models were downloaded from the pVOG database (<u>Grazziotin et al., 2017</u>) and merged into a single HMM database using *hmmpress*. For each species, predicted proteins were clustered using MMseqs2 with query and subject coverage of at least 60% and e-value of less than 0.001 (<u>Steinegger and Soding, 2017</u>). A representative from each cluster was searched against pVOG database using *hmmscan* (e-value < 1E-10, database size = 100,000) (<u>Eddy, 2011</u>). If the representative had a corresponding hit in the pVOG database, all proteins in the cluster were labelled as “phage-related proteins”.

To estimated the fraction of ‘phage-related proteins’ that are targeted by CRISPR-Cas systems, nucleotide sequences of phage-related genes were extracted from the FASTA file. The spacers were used as queries in an MMseqs2 search against phage-related genes with the minimum sequence identity of 95% and the minimum spacer coverage of 90%.

### Phylogenetic profiling of defense systems

To minimize spurious associations due to the phylogenetic signal, for each orthologous group, the Fritz and Purvis’ D-value (with 1,000 permutations) was calculated using the caper package (<u>Fritz and Purvis, 2010</u>; <u>Orme et al., 2013</u>). Only orthologous groups with D-values greater than –0.2 were retained for subsequent analyses. Furthermore, orthologous groups that were present in less than 10% of the genomes or more than 90% of the genomes were discarded. Phylogenetic profiling was conducted using Coinfinder, and p-values were adjusted via the Bonferroni correction (<u>Whelan et al., 2020</u>). Only defense systems with D-values greater than – 0.2, were retained. The orthologous group was considered associated with the multigene defense system if it showed a significant association with all genes of this defense system. The significantly associated orthologous groups were annotated using the CDD database v3.2 (<u>Lu et al., 2020</u>), and BLASTP search against the UniProtKB database (e-value < 0.1) (<u>Altschul et al., 1990</u>; <u>UniProt Consortium, 2021</u>).

## Author contributions

RK initiated the study and collected the data; RK, YIW and EVK analyzed the data; RK and EVK wrote the manuscript that was edited and approved by all authors.

## Acknowledgements

The authors’ research is funded by the Intramural Research Program of the National Institutes of Health (National Library of Medicine).

## Data availability statement

Derived data and custom scripts are deposited at Zenodo (DOI: 10.5281/zenodo.10637202). The IDs of the analyzed assemblies are provided in Supplementary Table 1 and publicly available in the RefSeq database.

