## Supplementary figures for "Defense systems and horizontal gene transfer in bacteria"

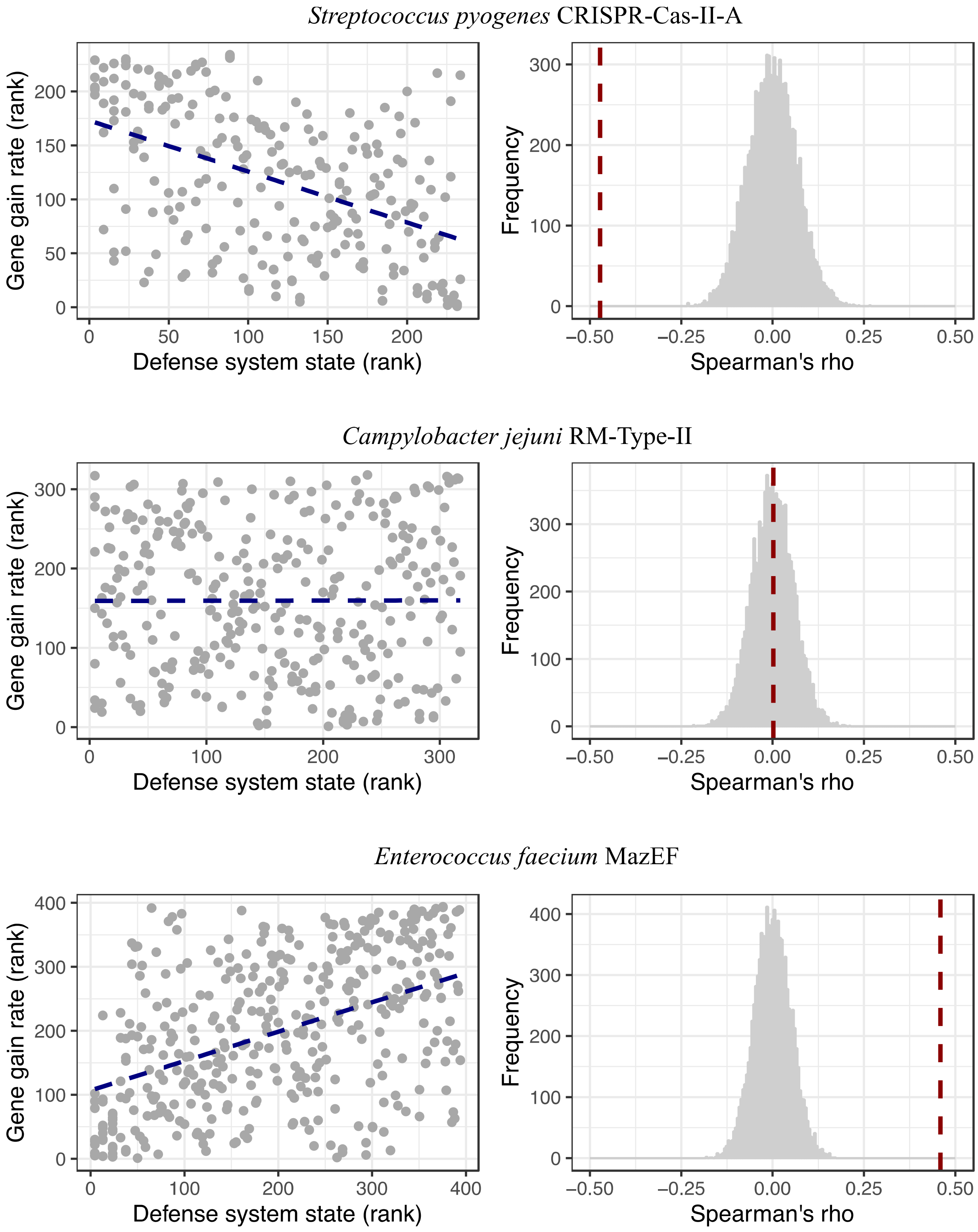


**Figure S1**. **Three defense systems that exhibit different correlation types with a gene gain rate.** On the scatter plot, each dot represents a branch on the phylogenetic tree, and a blue line shows the linear fit between axes. Accompanying histograms illustrate the null distribution of Spearman's rho values (in gray), while the observed Spearman's rho is highlighted by the dashed red line.

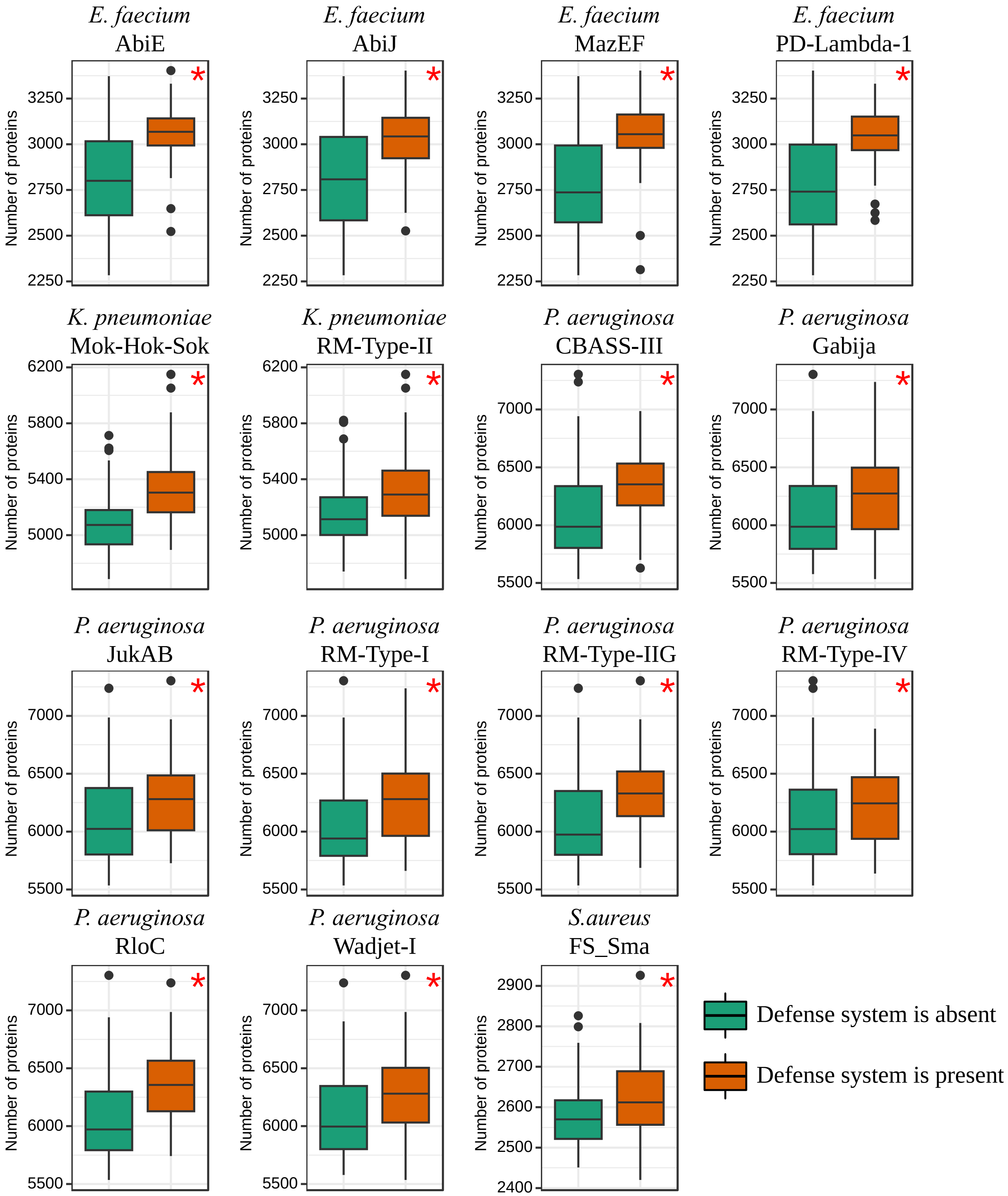


**Figure S2**. **Relationship between the 15 defense systems associated with the increased gene gain rate and number of protein coding genes in genomes.** Boxplots show median values that are bound by the first and third quartiles. Whiskers extended outside boxplots are within the 1.5

* interquartile range. Black dots outside whiskers denote outliers. Red asterisks show significant differences between boxplots (Mann-Whitney U test, p-value < 0.05).

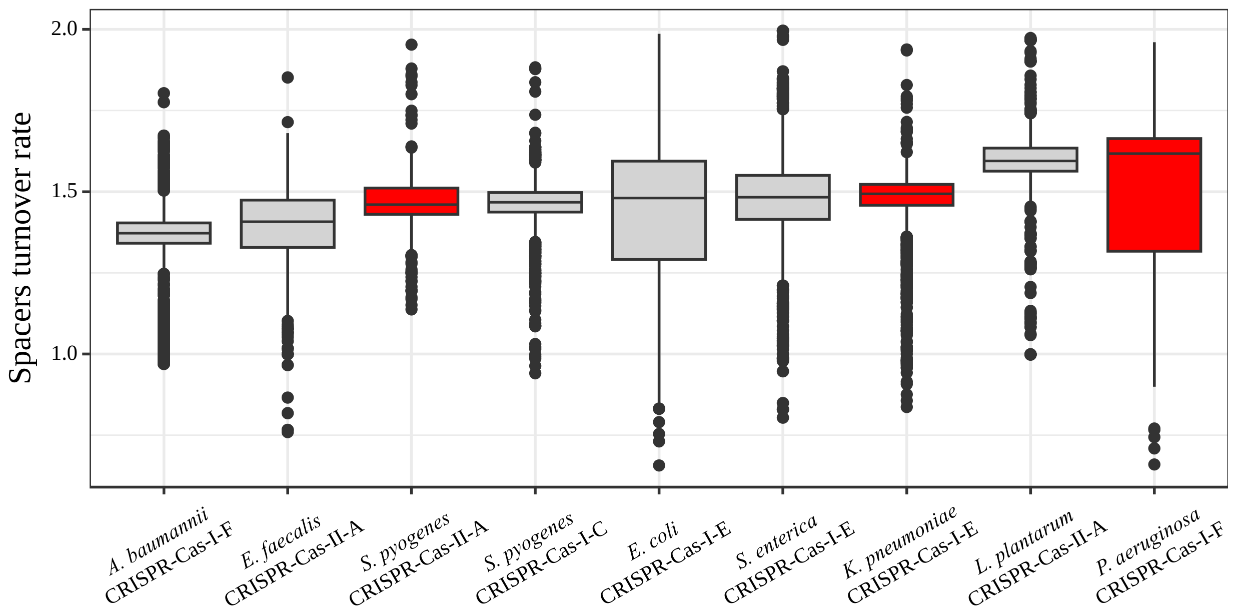


**Figure S3**. **Spacer turnover rate in 9 analyzed CRISPR-Cas systems.** Boxplots show median values that are bound by the first and third quartiles. Whiskers extended outside boxplots are within the 1.5 * interquartile range. Black dots outside whiskers denote outliers. Boxplots were sorted in the ascending order by their medians. Red boxplots red denote CRISPR-Cas systems associated with a reduced gene gain rate.

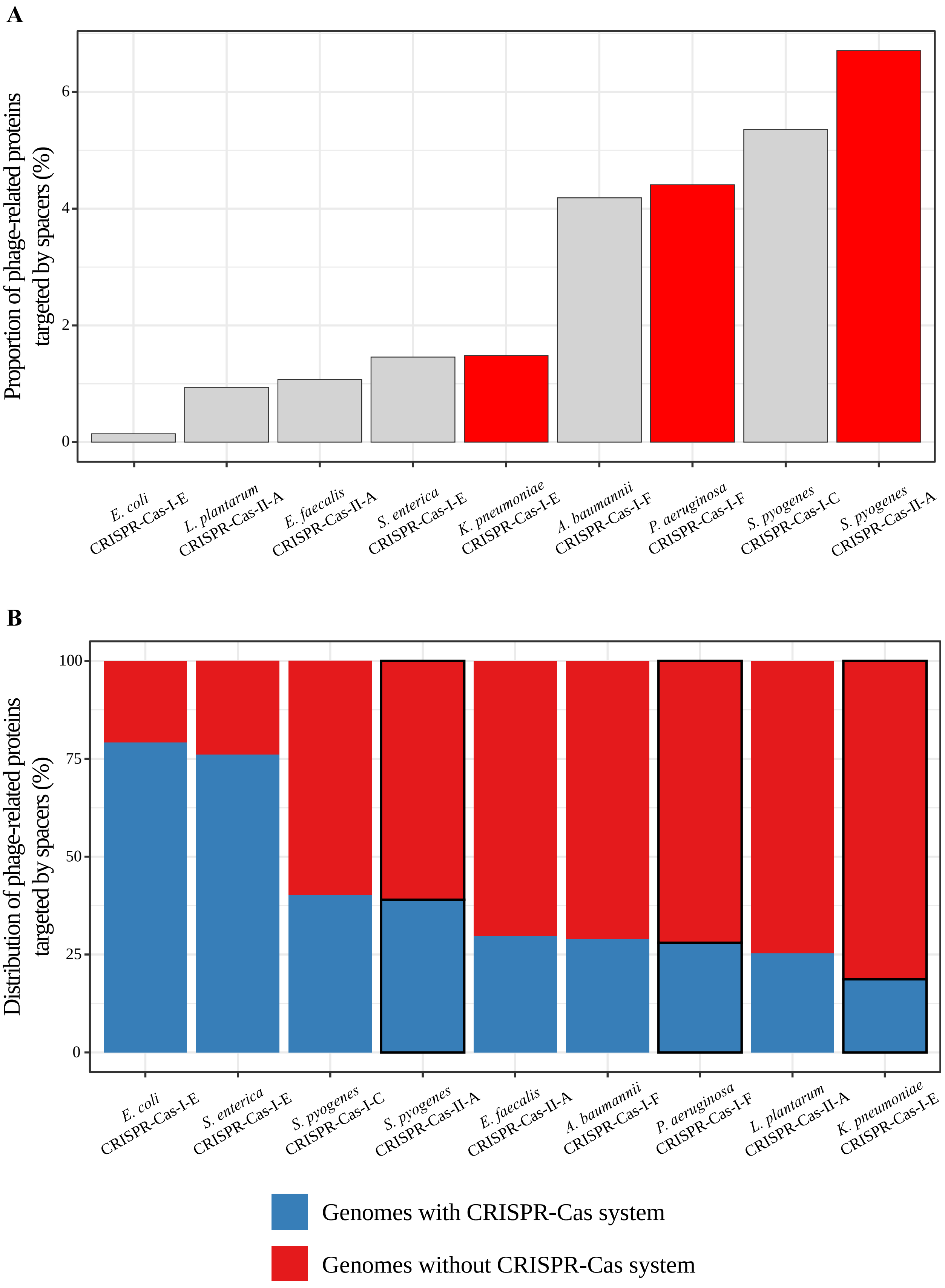
A
Proportion of phage-related proteins targeted by spacers (%)
0
2
4
6
E. coli CRISPR-Cas-I-E
L. plantarum CRISPR-Cas-II-A
E. faecalis CRISPR-Cas-II-A
S. enterica CRISPR-Cas-I-E
K. pneumoniae CRISPR-Cas-I-E
A. baumannii CRISPR-Cas-I-F
P. aeruginosa CRISPR-Cas-I-F
S. pyogenes CRISPR-Cas-I-C
S. pyogenes CRISPR-Cas-II-A
B
Distribution of phage-related proteins targeted by spacers (%)
0
25
50
75
100
E. coli CRISPR-Cas-I-E
S. enterica CRISPR-Cas-I-E
S. pyogenes CRISPR-Cas-I-C
S. pyogenes CRISPR-Cas-II-A
E. faecalis CRISPR-Cas-II-A
A. baumannii CRISPR-Cas-I-F
P. aeruginosa CRISPR-Cas-I-F
L. plantarum CRISPR-Cas-II-A
K. pneumoniae CRISPR-Cas-I-E
Genomes with CRISPR-Cas system
Genomes without CRISPR-Cas system

**Figure S4**. **Analysis of spacer composition in 9 CRISPR-Cas systems. (A)** Proportion of pooled phage-related genes that are targeted by CRISPR-Cas systems. Bar plots colored by red denote CRISPR-Cas systems associated with a reduced gene gain rate. **(B)** Distribution of phage-related genes that are targeted by spacers in genomes with and without CRISPR-Cas systems. CRISPR-Cas systems associated with the reduced gene gain rate are highlighted with the black rectangle.

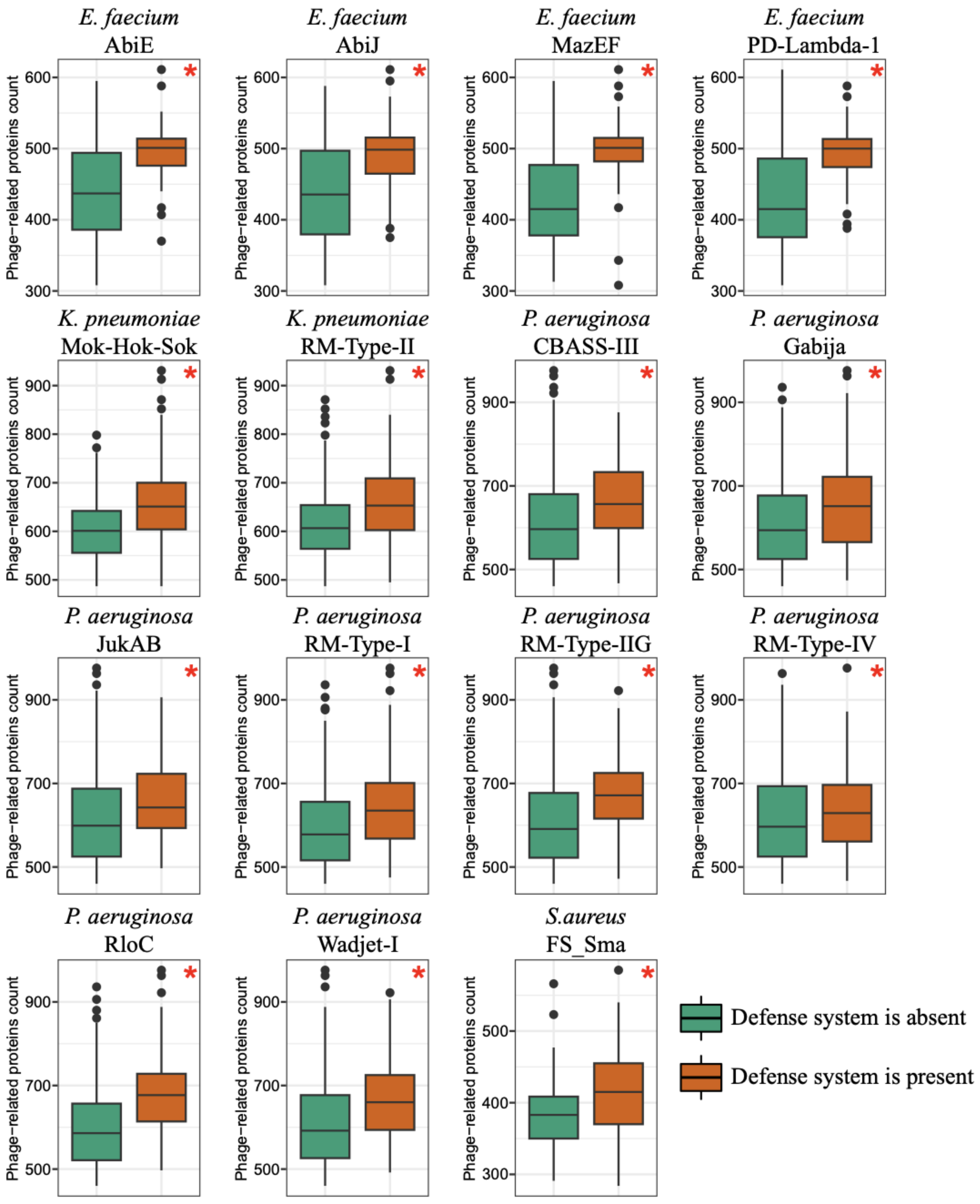


**Figure S5. Relationship between the 15 defense systems associated with an increased gene gain rate and the number of phage-related genes in genomes.** Boxplots show median values that are bound by the first and third quartiles. Whiskers extended outside boxplots are within the 1.5 * interquartile range. Black dots outside whiskers denote outliers. Red asterisks show significant differences between boxplots (Mann-Whitney U test, $p\text{-value} < 0.05$).
